# *De novo* Identification of Essential Protein Domains from CRISPR/Cas9 Tiling-sgRNA Knockout Screens

**DOI:** 10.1101/581751

**Authors:** Wei He, Liang Zhang, Oscar D. Villarreal, Rongjie Fu, Ella Bedford, Jingzhuang Dou, Mark T. Bedford, Xiaobing Shi, Taiping Chen, Blaine Bartholomew, Han Xu

**Affiliations:** Department of Epigenetics and Molecular Carcinogenesis, The University of Texas MD Anderson Cancer Center, Smithville, Texas, 78957, USA.; The Center for Cancer Epigenetics, The University of Texas MD Anderson Cancer Center, Houston, Texas 77030, USA.; Center for Epigenetics, Van Andel Research Institute, Grand Rapids, Michigan 49503, USA.; Department of Bioinformatics and Computational Biology, The University of Texas MD Anderson Cancer Center, Houston, Texas 77030, USA.

## Abstract

High-throughput CRISPR/Cas9 knockout screens using a tiling-sgRNA design permit *in situ* evaluation of protein domain function. To facilitate *de novo* identification of essential protein domains from such screens, we developed ProTiler, a computational method for the robust mapping of CRISPR knockout hyper-sensitive (CKHS) regions, which refers to the protein regions that are associated with strong sgRNA dropout effect in the screens. We used ProTiler to analyze a published CRISPR tiling screen dataset, and identified 175 CKHS regions in 83 proteins. Of these CKHS regions, more than 80% overlapped with annotated Pfam domains, including all of the 15 known drug targets in the dataset. ProTiler also revealed unannotated essential domains, including the N-terminus of the SWI/SNF subunit SMARCB1, which we validated experimentally. Surprisingly, the CKHS regions were negatively correlated with phosphorylation and acetylation sites, suggesting that protein domains and post-translational modification sites have distinct sensitivities to CRISPR/Cas9 mediated amino acids loss.

## Introduction

Functional screens using CRISPR/Cas9 techniques facilitates the identification of essential genes on a genome-wide scale^1–4^. To fully define the function of protein-coding essential genes, it is necessary to distinguish essential protein domains that directly contribute to cellular phenotypes. In a CRISPR/Cas9 knockout experiment, the sgRNAs that target DNA sequences coding for essential protein domains often result in more significant dropout phenotype compared to other sgRNAs that target the same gene^5^. This is likely because CRISPR/Cas9 introduces small indels that create frameshift or in-frame mutations in a stochastic manner. Frameshift indels tend to abolish protein function, whereas in-frame indels, which result in the gain or loss of amino acids, may or may not impact function depending on where they occur. Proteins with small, in-frame indels in nonessential regions are likely to retain function. In contrast, proteins with indels in essential domains may display compromised protein function due to the disruption of an important structural motif or functional conformation (Fig. 1a). Therefore, a domain-focused CRISPR/Cas9 knockout screen has been proposed to evaluate the functional importance of individual domains, leading the way for *in situ* protein functional studies^5^.

**Fig. 1.**
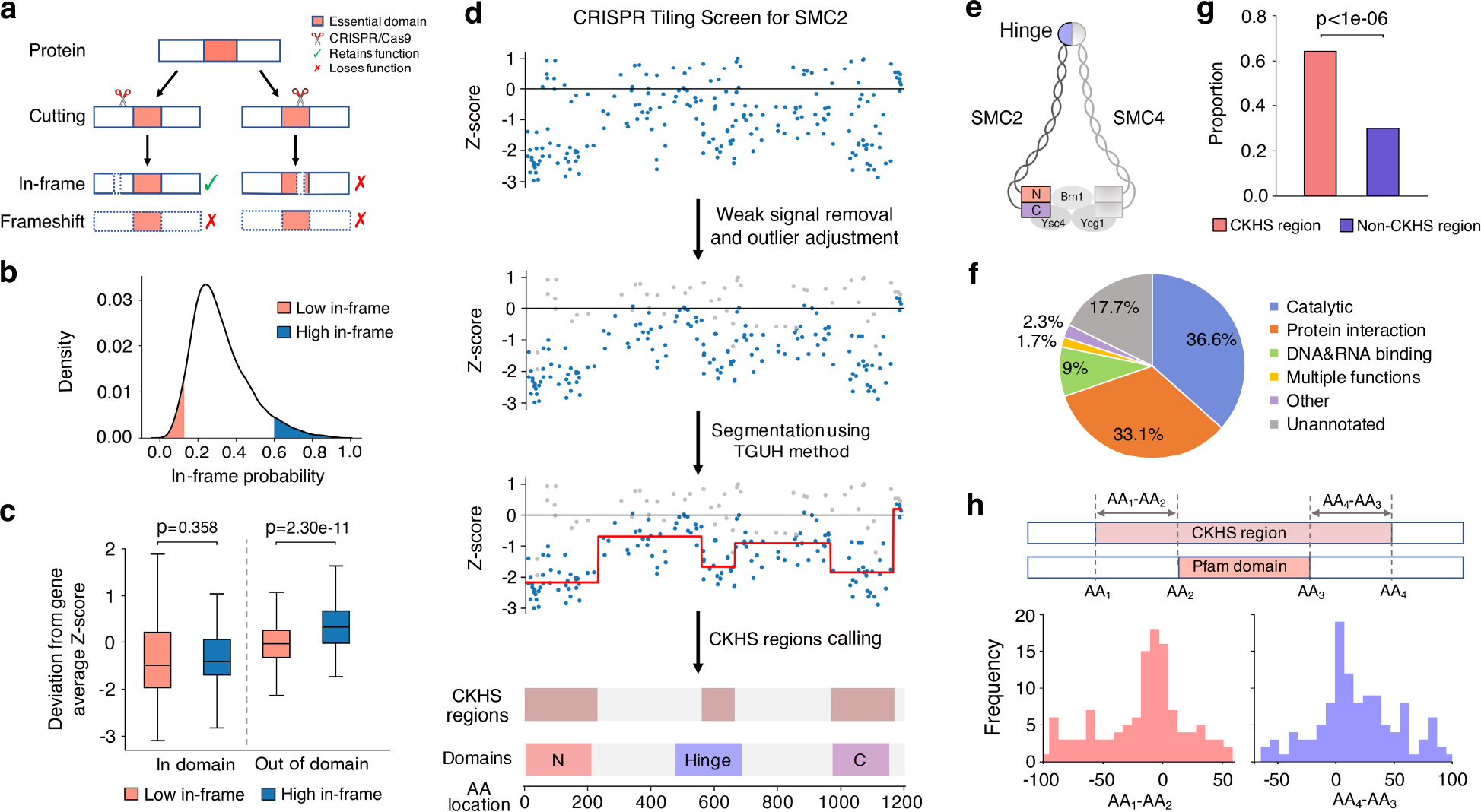
Mapping CRISPR knockout hyper-sensitive (CKHS) regions with ProTiler. **a)** An in-frame indel model underlying the rationale of domain-associated CKHS regions. **b)** Distribution of in-frame probability of the sgRNAs targeting essential proteins in Munoz data. The probabilities were predicted using inDelphi^11^. The Top 5% and bottom 5% sgRNAs are defined as “high in-frame” (blue) and “low in-frame” (red), respectively. **c)** Box-plots comparing the dropout effects between “high in-frame” and “low in-frame” sgRNAs that target proteins containing drug target domains. The p-values were computed using the Mann-Whitney test. **d)** The workflow of ProTiler. The dot plots show the dropout effects, in Z-score^6^ (Y-axis), of sgRNAs targeting the gene coding for SMC2. A negative value of the Z-score corresponds to a dropout effect. Each dot represents an sgRNA mapped to the amino acid location (X-axis). The grey dots and blue dots represent filtered and remaining sgRNAs, respectively. The red line shows the segmented protein regions and their dropout signal levels. **e)** A structural model of condensin complex, in which SMC2 and SMC4 form a heterodimer via hinge domains, and their ATPase head domains (N and C) are associated with kleisin subunits to create a ring-like structure. **f)** Categorization of CKHS regions based on the molecular functions of overlapped protein domains. **g)** A bar chart showing the proportion of AAs in Pfam domains, for CKHS regions and non-CKHS regions respectively. The p-value was empirically computed by random simulation. **h)** Distribution of distances between the borders of CKHS regions and domain boundaries as defined in the Pfam database.

In a pooled high-throughput CRISPR/Cas9 knockout screen, an sgRNA library contains tens of thousands of sgRNAs. This depth of coverage facilitates a tiling-sgRNA design that allows the investigation of domain functions across the entire protein for more than 100 protein-coding genes in a single experiment. Munoz *et al.* performed the first high-throughput tilling-sgRNA screen on 159 genes, and confirmed that the sgRNAs that target pharmaceutically important protein domains are associated with stronger knockout effects^6^. Recently, a method combining tiling-sgRNA screens with positive selections has been developed to identify small-molecule drug target sites^7^. A computational pipeline, CRISPRO, maps functional scores of tiling sgRNAs to genomes, transcripts, protein coordinates and structures, providing general views of structure-function relationships at discrete protein regions^8^.

Despite these advances, pooled high-throughput CRISPR/Cas9 screens are subject to inactive sgRNAs, off-target effects, and high noise-to-signal ratios, posing computational challenges to robust identification of essential domains. To address these challenges, we developed ProTiler, a computational method for the mapping of protein regions that are associated with strong sgRNA dropout effect in the screens, termed CRISPR knockout hyper-sensitive (CKHS) regions. We applied ProTiler to the dataset published by Munoz *et al.*, aiming at a systematic evaluation of tiling-sgRNA screens for *in situ* protein functional analysis, as well as the potential for novel domain discovery.

## Results

### Essential protein domains are hyper-sensitive to CRISPR/Cas9 induced in-frame indels

The basis of using tiling-sgRNA screens for protein domain analysis relies on an in-frame indel model, as shown in Fig. 1a. Recently, several laboratories showed that CRISPR/Cas9 indel patterns and in-frame mutational probabilities are predictable from sgRNA-targeted DNA sequences^9, 10^. Taking advantage of these findings, we first examined if the model is applicable to Munoz data. We used inDelphi^11^ to measure the in-frame probabilities for 28,951 sgRNAs, corresponding to 108 essential protein-coding genes. The median in-frame probability is 0.29, suggesting incomplete protein knockout in approximately half of diploid cells. To ensure robustness against prediction error, we selected the sgRNAs that were predicted to be very likely (top 5%, in-frame probability > 0.60, “high in-frame”) or very unlikely (bottom 5%, in-frame probability < 0.129, “low in-frame”) to create in-frame mutations (Fig. 1b). Consistent with the model, the “low in-frame” sgRNAs showed greater dropout effect compared to the “high in-frame” sgRNAs (p=4.85e-76, Supplementary Fig. 1). To assess whether the difference between two categories is associated with the functional essentiality of protein domains, we examined 15 domains targeted by small-molecule compounds that have either been FDA-approved or advanced to clinical trials (Supplementary Table 1). Among the sgRNAs that target DNA sequences coding for these domains, no significant difference was observed between the “high in-frame” and the “low in-frame” categories (p=0.358). In contrast, sgRNAs associated with the regions outside annotated domains showed significant difference (p=2.30e-11, Fig. 1c). These results indicate that the essential domains are hyper-sensitive to CRISPR/Cas9 induced in-frame indel mutations, supporting the underlying rationale of using tiling-sgRNA screens to predict domain essentiality.

### ProTiler enables fine-mapping of protein regions hyper-sensitive to CRISPR/Cas9 knockouts

We developed ProTiler, a computational method for the mapping of protein regions that are associated with CRISPR knockout hyper-sensitivity (CKHS). Fig. 1d outlines the major steps in ProTiler, exemplified by tiling-sgRNA screen data for SMC2, a component of the condensin complex. ProTiler first maps sgRNA dropout signals to the amino acids of the target proteins. The data points with weaker dropout effects compared to their neighbors are likely to be associated with inactive sgRNAs (Supplementary Fig. 2), thus are removed. The “outliers” in the remaining data points can be caused by non-Gaussian variations or additive off-target effects. We adjusted their values based on the mean and variation of the surrounding signals. To partition the protein into regions corresponding to different viabilities, we applied Tail-Greedy Unbalanced Haar (TGUH) transformation, a wavelet-based changing point detection algorithm with proven high accuracy and robustness under noisy conditions^11^. Each region is assigned a viability score to be the average of the data points in that region. Finally, an iterative algorithm classifies the regions into CKHS and non-CKHS categories. For SMC2, ProTiler detected three CKHS regions, corresponding to the N-terminus, C-terminus and the middle hinge domain. These CKHS regions are highly consistent with a model of condensin structure, in which SMC2 and SMC4 form a heterodimer via their hinge domains and their ATPase head domains are associated with kleisin subunits to create a ring-like structure^12^ (Fig. 1e).

Among 108 essential proteins in the Munoz data, ProTiler identified 175 CKHS regions in 83 proteins. 82.3% of these regions overlapped with Pfam annotated protein domains (Fig. 1f, Supplementary Table 3). At the amino acid (AA) level, 64.2% of the AAs in the CKHS regions are within Pfam domains, compared to 30.0% for non-CKHS regions (Fig. 1g). All of the 15 previously mentioned drug target domains were identified (Supplementary Table 1), suggesting high sensitivity. To estimate the resolution, we aligned the borders of ProTiler-defined CKHS regions to the boundaries of Pfam domains. The borders CKHS regions were enriched within 20 AAs from domain boundaries and slightly outside the domains, suggesting that small deletions of AAs adjacent to the domains may also compromise protein function (Fig. 1h). Taken together, these lines of evidence indicate high performance of ProTiler, in specificity, sensitivity, and resolution.

### CKHS mapping facilitates discovery of novel essential domains

Among the ProTiler-identified CKHS regions, 17.7% did not overlap with any annotated Pfam domains, which may be associated with novel or previously undefined domains. Indeed, some of these regions have been functionally characterized but remain unannotated in the Pfam database. For example, a small CKHS region in the transcriptional coactivator YAP1 (AA 86-99) perfectly matches a twisted-coil structure, one of the three independent interfaces that interacts with TEAD1 (Fig. 2a). Consistent with previous findings, our results showed the importance of YAP1-TEAD1 interaction for cell viability, where the region of AA 86-99 is the most critical interaction site^13^ (Fig. 2b). ProTiler also identified a CKHS region in the N-terminus (AA 55-184) of MPS1/TTK, which contains three tetratricopeptide repeat domains that govern localization of the protein to either the kinetochore or the centrosome^14^ (Supplementary Fig. 3)

**Fig. 2.**
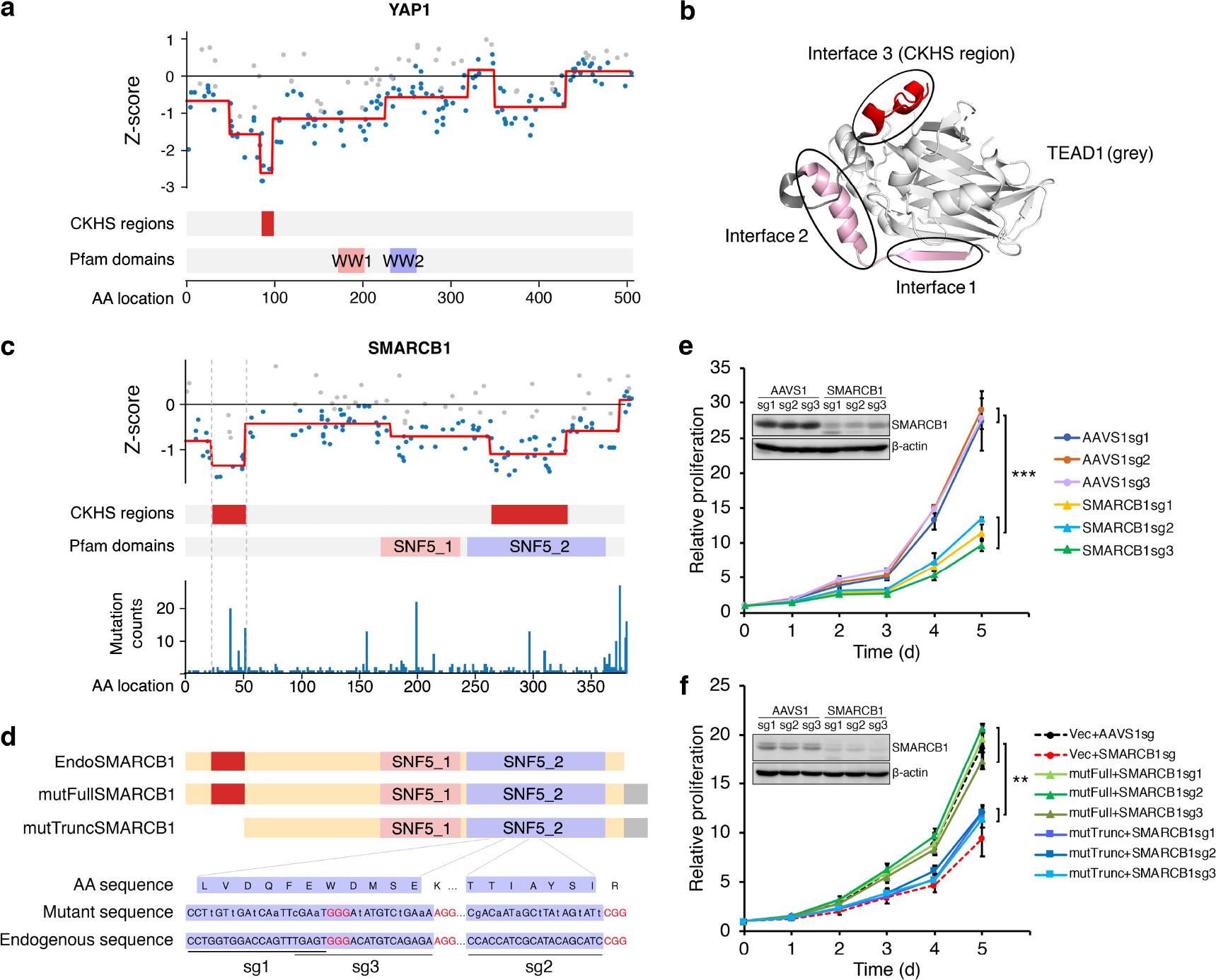
CKHS profiling facilitates the identification of unannotated essential domains. **a)** The CKHS profile and domain annotation of YAP1. **b)** The 3D structure of the YAP1&TEAD1 interaction (PDBID: 3KYS). Three interfaces of YAP1 interacts with TEAD1. The CKHS region (interface 3) is highlighted in red. **c)** The CKHS profile and domain annotation of SMARCB1, aligned with mutation frequency retrieved from the COSMIC database^46^. **d)** A schematic representation of exogenous full-length or a truncated form (ΔAA 2-53) of SMARCB1, as well as the endogenous protein. The CKHS region is highlighted in red. A Myc and 6xHis tag sequence were appended to the C-terminus of exogenous proteins. Three sgRNAs were designed to target the endogenous *SMARCB1* DNA sequence coding for the SNF5_2 domain. Synonymous mutations were introduced into the exogenous proteins, except the bases at the PAM (red) and methionine codon. **e)** Proliferation of DLD-1 cells after SMARCB1 knockout by CRISPR/Cas9. The sgRNAs targeting the *AAVS1* locus were used as the controls. All the data points represent the average of three biological replicates. ***: p<0.001, T-test. **f)** Proliferation of DLD-1 cells with exogenous expression of full-length or the truncated form of SMARCB1 shown in d), in combination of endogenous SMARCB1 knockout. The red and black dash lines represent proliferations of vector control cells with or without SMARCB1 knockout, respectively. All the data points represent the average of three biological replicates. **: p<0.01. T-test.

Inspired by these examples, we sought to identify novel essential domains within newly identified, but unannotated CKHS regions. After careful inspection, we focused on SMARCB1, one of the core subunits of the SWI/SNF chromatin remodeling complex, for further validation. ProTiler identified a CKHS region (AA 24-53) in the N-terminus of SMARCB1, in addition to the well-characterized ATP-binding domain. We chose to examine this region because it habors recurrent missense mutations (P48L, R53L, E31L) and frameshift deletions (G29, S30) that have been identified in cancer patients^15^ (Fig. 2c). An earlier *in vitro* study showed that the N-terminus of SMARCB1 forms a putative DNA-binding structure^16^, but its function has not been investigated *in situ*. To validate the function of this region, we constructed vectors expressing either a full-length or an N-terminally-truncated form of SMARCB1 (Fig. 2d, Supplementary Fig. 4). In the constructs, we introduced synonymous mutations to the ATP-binding domain, such that the knockouts with sgRNAs targeting the mutated sites would affect only the endogenous SMARCB1 without compromising the expression of exogenous proteins. The vectors were lenti-viral introduced into DLD-1, a colon cancer line expressing a wild-type SMARCB1. Consistent with the screen data, CRISPR-mediated knockouts of SMARCB1 hampered the growth of DLD-1 cells (Fig. 2e), which could be completely rescued by exogenous expression of the full-length protein, but not the truncated protein (Fig. 2f). Collectively, these lines of evidence confirmed the essential role of the CKHS region in the N-terminus of SMARCB1.

### CKHS profiling of multi-domain proteins reinforces recent findings in the literature

Proteins containing multiple functional domains often have complex cellular roles, thereby posing challenges to their molecular characterization. A tiling-sgRNA screen can facilitate the assessment of domain functions in a high-throughput manner. We explored the CKHS profiles of 51 multi-domain proteins in the Mounz dataset. Domains in these proteins are associated with a wide range of functions, including catalysis, protein-protein interactions, and DNA/RNA binding. For these 51 proteins, 62.1% of the domains were marked with CKHS regions (Supplementary Table 4). Catalytic domains had the highest likelihood (93.2%) of being essential, compared to the others (55.0%, p=3.71e-07, Fig. 3a). Although it is beyond the scope of this study to elucidate the mechanism underlying the domain essentiality for each protein, we sought to link CKHS profiles to recent findings in the literature, as a proof of value for novel protein functional discovery. Here, we present three examples.

**Fig. 3.**
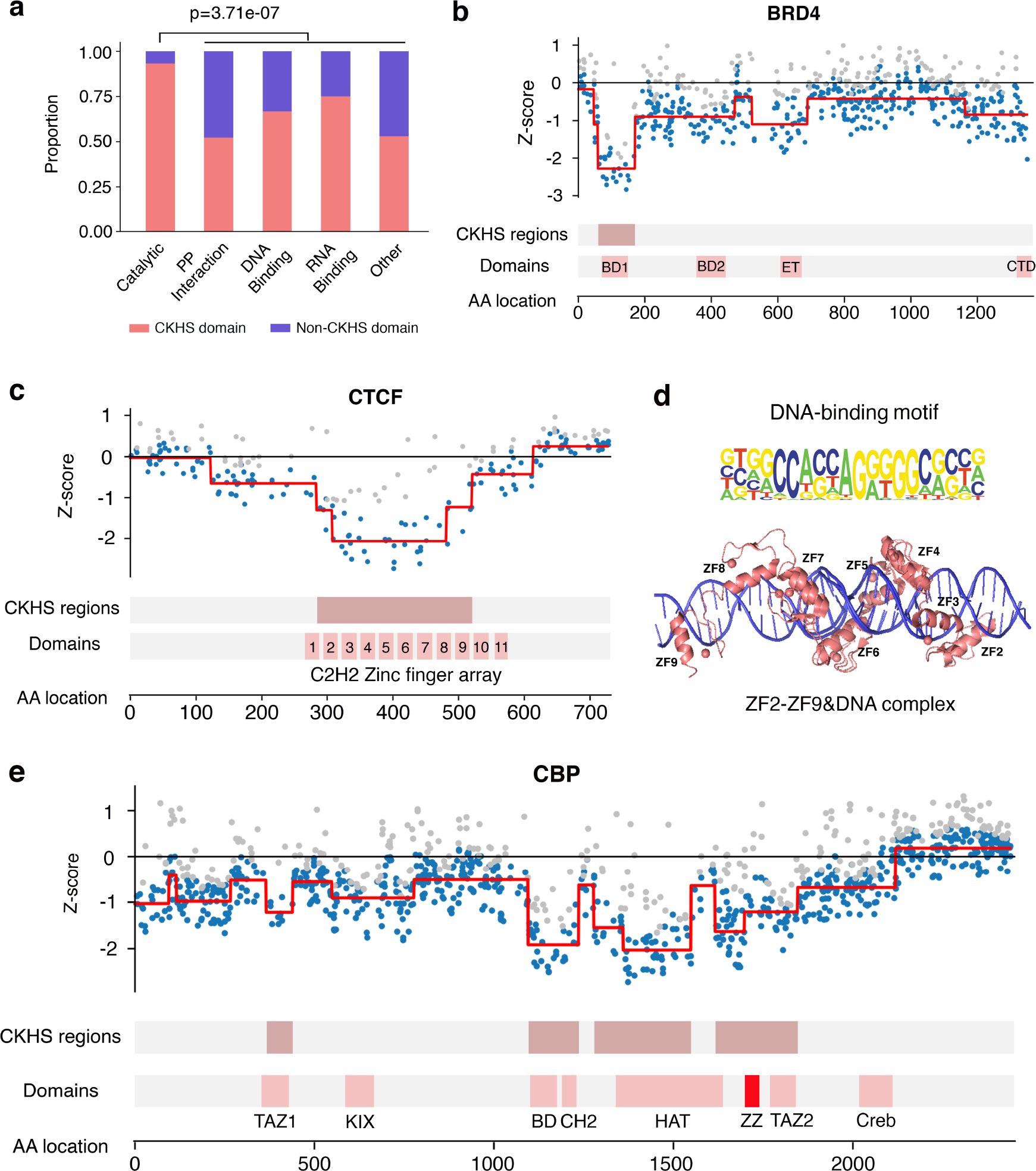
The CKHS profiles of multi-domain proteins. **a)** All the domains in 51 multi-domain proteins were categorized based on their molecular functions. The bar chart shows the proportion of CKHS-overlapped domains for each category. The p-value was computed based on hyper-geometric distribution. **b)** The CKHS profile and domain annotation of BRD4. **c)** The CKHS profile and domain annotation of CTCF. **d)** The 3D structure of CTCF (ZF2-ZF9) complexed with DNA (PDB IDs: 5UND;5T0U). The DNA binding motif ^47^ is aligned with the structure. **e)** The CKHS profile and domain annotation of CBP. The ZZ domain is highlighted in red

BRD4 is an acetyl-lysine reader known for its function as a transcriptional co-activator^17^. It is also a major target of several BET inhibitors that have progressed to clinical trials^18^. BRD4 contains two bromodomains (BD1 and BD2), an extra-terminal domain (ET), and a C-terminal domain(CTD). Despite high sequence similarity between BD1 and BD2, only BD1 showed hyper-sensitivity to CRISPR knockouts (Fig. 3b). In line with this observation, an earlier investigation reported that inhibition of BD2 inhibition had a milder effect on BRD4-dependent gene transcription than BD1 inhibition^19^. Recently, several laboratories reported a BD1-dependent role of BRD4 in the DNA damage response pathway, raising additional concerns regarding BRD4 function involved in the viabilities of normal and cancer cells^20–22^. To this end, our results highlighted the importance of bromodomain selectivity in the functional analysis and drug discovery regarding BRD4 inhibition.

CTCF is a DNA-binding protein critical for maintaining high-order chromatin conformations^23^. It has 11 adjacent zinc-finger domains (ZF1-ZF11) that bind to DNA. Its CKHS profile showed that only ZF2-ZF9 are hyper-sensitive, suggesting that individual ZFs have unequal contributions to CTCF function (Fig. 3c). Recently, a crystal structure of CTCF-DNA interaction was released, which showed that ZF2-ZF9 are required for the binding to the full nucleotide motif sequence^24^ (Fig. 3d). Thus, our high-resolution CKHS map is supported by this structure model, and in turn reinforces the new structural model *in situ*.

CBP, a paralog of p300, is a transcriptional co-activator that has been extensively studied over the last two decades^25^. It has a catalytic core containing a HAT domain, a bromodomain, and a CH2 region^26^. All the three regions were found to be CKHS in our results (Fig. 3e). Two protein-interacting domains, TAZ1 and TAZ2, were also sensitive to CRISPR knockouts but to a lesser degree. In addition to these well-characterized domains, a ZZ-type zinc finger domain adjacent to the HAT domain is also in the CKHS region. Recently, the ZZ domain was characterized to be an acetyl-reader of histone H3, which modulates CBP/p300 enzymatic activity and their associations with chromatin^27^. Therefore, the CKHS profile of CBP further supports the critical role of the ZZ domain.

Taken together, these examples indicate that the CKHS profiles can be used either to infer a new functional model, or to validate an existing model. Potential applications include, but are not limited to, prediction of potent inhibitor targets, discovery of alternative protein functions, *in situ* validation of protein structure, and identification of novel domain function.

### CKHS regions are negatively correlated with phosphorylation and acetylation sites

Despite the consistency between CKHS regions and essential domains in CBP, we observed that a small region in the HAT domain (AA1550-1618) was insensitive to CRISPR knockouts. This region matches the auto-acetylation sites that are known to regulate the catalytic activity of CBP^28^ (Fig. 4a). Similarly, a cluster of phosphorylation sites between the kinase domain and the RAS-binding domain of BRAF also showed hypo-sensitivity despite of their critical role in regulating BRAF activity^29^. Another group of phosphorylation sites near AA600 of BRAF is in the CKHS region, but showed lesser sensitivity compared to the adjacent kinase domain (Fig. 4b). Indeed, when we mapped the post-translational modification (PTM) sites onto our data, we found phosphorylation and acetylation sites were significantly depleted inside CKHS regions compared to outside CKHS regions (Fig. 4c). Therefore, the sensitivity to CRISPR knockouts is negatively correlated with phosphorylation and acetylation sites. Since those PTMs are often clustered to modulate protein conformation via electric charges, a possible explanation to this observation is that, a CRISPR/Cas9 mediated small deletion of amino acids near PTMs does not significantly reduce charges in a local region, which in turn has weaker impact on protein conformation and phenotype. Additionally, methylation and ubiquitination are also statistically associated with CKHS regions, but the explanations for these associations are yet elusive. Collectively, these observations indicate that protein domains and PTM sites have distinct sensitivities to CRISPR/Cas9 mediated amino acids loss, thus a careful inspection on PTM annotations will be helpful for the interpretation of tiling-sgRNA screen data.

**Fig. 4.**
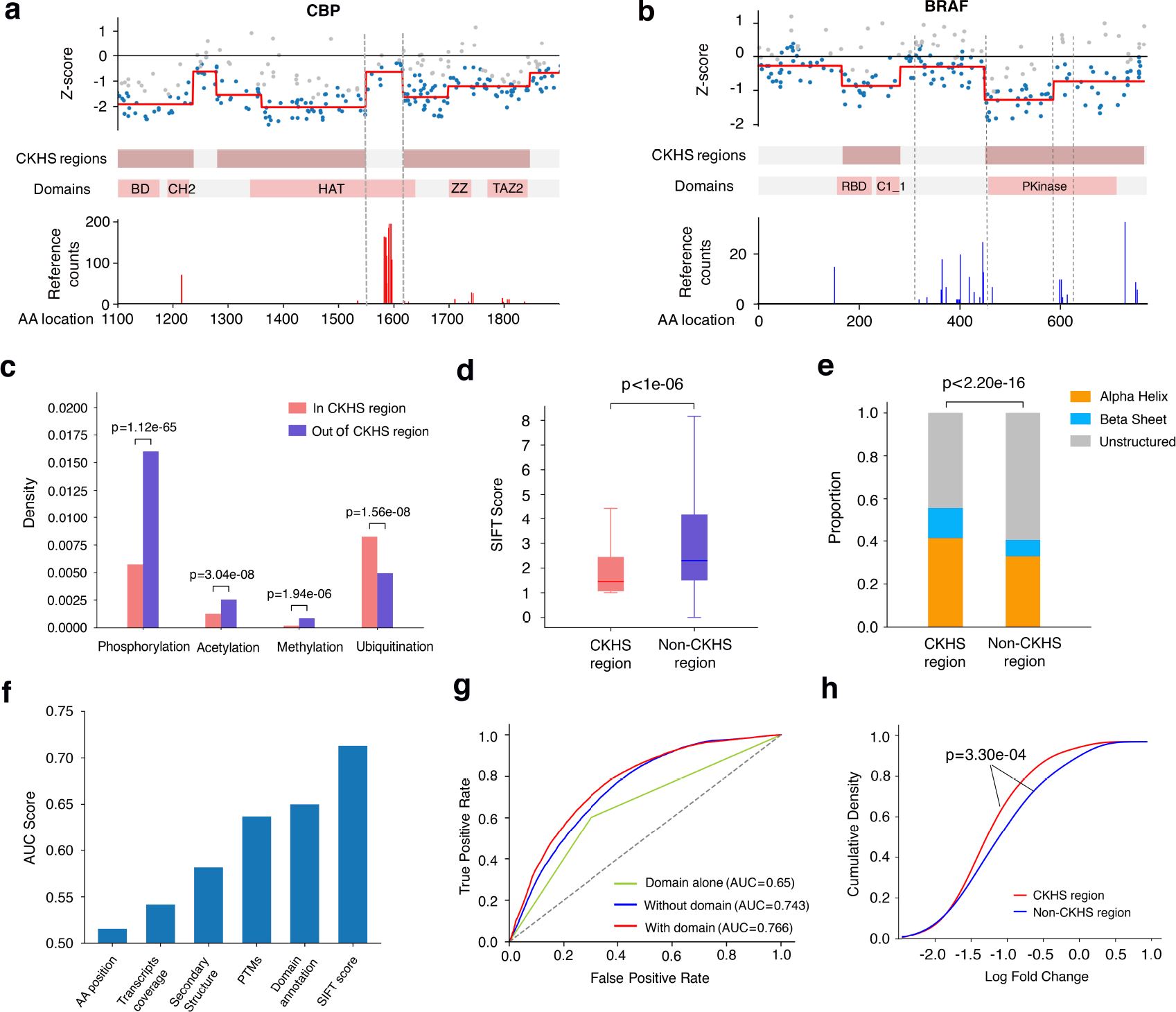
CKHS regions are associated with post-translational modifications (PTMs) and are predictable from protein features. **a)** A zoomed-in views of CKHS profile, domain annotation, and acetylation sites of CBP (AA 1100-1900). The Y-axis of the PTM profiles represent the number of publication references collected at https://www.phosphosite.org. **b)** The CKHS profile, domain annotation, and phosphorylation sites of BRAF. **c)** A bar chart showing the density of PTM sites inside or outside of the CKHS regions. The p-values were computed based on hypergeometric distribution. **d)** A box-plot showing the association between CKHS regions and amino acids conservation (SIFT score). The p-value was computed using the Mann-Whitney test. **e)** A bar chart showing the distribution of secondary structures for CKHS and non-CKHS regions. The p-value was computed using the Chi-square test. **f)** A bar chart showing the predictive power of bagging SVM, in ROC-AUC score, using individual protein features. The AA position in the protein and the transcript coverage are used as references. **g)** ROC curves showing the predictive powers using all protein features, all features other than domain annotation, and domain annotation alone, respectively. **h)** The sgRNAs targeting core essential genes were categorized based on predicted CKHS regions. The cumulative distributions of sgRNA dropout effects in Avana dataset^35^ are shown for each category. The p-values were computed using Kolmogrov-Smirnov test.

### A proteome-wide prediction of CKHS regions for the rational design of CRISPR libraries

The sgRNA knockout effects are associated with both the level of amino acid sequence conservation and the secondary structure of target protein regions^8^. CRISPRO introduced a machine-learning approach for the prediction of CRISPR/Cas9 knockout effect, using the features related to protein function and sequence-specific sgRNA activity. Consistently, we found that CKHS regions are associated with highly conserved regions (SIFT score) and molecular secondary structures (Fig. 4d, e). Since the tiling screen data are available for only a limited number of proteins, we sought to predict CKHS regions based on protein features in a proteome-wide scale. Recently, several variants of the CRISPR system have been developed for genome editing^30–32^, each associated with a unique sequence preference for sgRNA activity. Therefore, we aimed at a predictive model based on protein features alone, such that the predicted CKHS regions would be independent of the sgRNA target sequences and could be used for different CRISPR techniques.

Using a bagging Support Vector Machine (bagging SVM)^33^ and a “leave-one-gene-out” cross-validation strategy, we first examined the predictive power of individual features. The SIFT score showed the highest power (AUC=0.713), followed by domain (AUC=0.650) and PTM (AUC=0.632) annotations (Fig. 4f). As references, transcript variant coverage and AA position in relative to protein N- and C-termini have little predictive power, indicating that protein functional features are the determinants of CKHS profile. Our model further achieved a higher performance when it integrated SIFT score, secondary structure, protein domain and PTM annotations (Fig. 4g). Notably, excluding domain annotations did not significantly compromise the prediction, suggesting this approach can be applied to the proteins lacking domain information. A proteome-wide prediction of CKHS regions is available in Supplementary Table 5. Finally, we tested whether the predicted CKHS regions can improve the sgRNA design, using two large-scale CRISPR/Cas9 screen datasets^34, 35^. Our results showed that the selection of sgRNAs based on the predicted CKHS prediction achieved greater dropout effects for known essential genes, suggesting the potential of applying our model to the rational design of CRISPR sgRNA libraries (Fig. 4h and Supplementary Fig. 5).

## Discussion

Essential protein domains are often associated with hyper-sensitivity to CRISPR/Cas9 knockouts, permitting *in situ* analysis of protein functions. Previous applications have mainly focused on the validation of annotated or hypothetical domains^5, 7^. A pooled CRISPR screen allows a tiling-sgRNA design for more than 100 proteins, holding promise for *de novo* discovery of essential domains. Here, we propose ProTiler for the mapping of CKHS regions from a tiling-sgRNA CRISPR screen. The high performance of ProTiler enables the identification of multiple essential domains in a protein, exemplified by SMC2, CTCF and CBP in this article. Using ProTiler, we identified a novel essential domain in the N-terminus of SMARCB1, which was further experimentally validated. To the best of our knowledge, this is the first report of a *de novo* prediction of a novel essential domain from large-scale tiling-sgRNA screens. The CKHS regions are highly concordant with essential domains of different types, including those associated with catalysis, protein-protein interactions, and DNA or RNA binding. These results established the tiling-sgRNA approach as a general method for *in situ* protein functional studies. Of note, although this work is focused on identifying essential domains that contribute to cell viability, our approach can also be applied to the screens with other readouts or cell sorting systems to understand domain functions associated with various cellular phenotypes.

We showed that the observation of CKHS regions is a consequence of in-frame indel mutations that influce protein domain function (Fig. 1c). It is possible that some protein-independent factors, such as transcription variants, reverse transcription, enhancers, and RNA modification sites, also contribute to the knockout hyper-sensitivity. By mapping sgRNAs to transcript variants, we observed a weak but statistically significant association between the coverage of variants and the CKHS regions (Supplementary Fig. 6). On the other hand, neither the variant coverage nor AA position had sufficient power to predict the CKHS regions, suggesting protein domain function is the major determinant of CKHS regions (Fig. 4f). Nevertheless, it would be helpful to check other possible confounding factors when interpreting data arising from a tiling-sgRNA screen.

A typical design of a CRSIPR tiling sgRNA library includes all the target sequences followed by a PAM motif. Screens with such an “all-target” library are subject to inactive sgRNAs and off-target effects. ProTiler addresses these computational challenges by removing weak signals and adjustment of outliers. We note that a number of computational tools have been developed to predict on-target activities and off-targets^36–38^. These tools are useful for designing sgRNA libraries for genome-scale screens. However, filtering sgRNAs according to computational predictions will compromise the overall resolution of a tiling-sgRNA library, due to a considerable number of unavoidable computational false negatives. Similarly, sgRNA selection based on the prediction of in-frame probabilities may not be applicable for the design of tiling-sgRNA libraries, because only 11.2% of sgRNAs have a predicted in-frame probability greater than 0.5 (Fig. 1b). Therefore, an “all-target” or “nearly-all-target” library seems to be more plausible, where maximum information content is maintained. At the same time, it is likely that ProTiler can be further improved using an algorithm that prioritizes sgRNAs based on such computational predictions.

## Methods

### Datasets and external software

The Munoz dataset was retrieved from publication^6^. The Pfam domain annotation was downloaded from Pfam database^39^ (version 31.0). The PTM annotation was downloaded from PhosphoSitePlus database^40^. The Avana dataset and GeCKO dataset were downloaded from https://figshare.com/articles/DepMap_Achilles_19Q1_Public/7655150 and https://figshare.com/articles/DepMap_GeCKO_19Q1/7668407, respectively. The transcript annotation was downloaded from NCBI CCDS database^41^. The list of 15 drug targets were manually curated in reference to the publications in Supplementary Table 1. The in-frame probabilities were computed using inDelphi online version at https://indelphi.giffordlab.mit.edu/. The amino acid conservation scores were computed using SIFT^42^. The protein secondary structures were computed using RaptorX^43^.

### Identification of essential genes

The Munoz dataset includes tiling-sgRNA screens on three cell lines (RKO, NCIH1299, and DLD1). We computed the average Z-score for each gene in each cell lines. Using a threshold of - 0.4, as suggested in the original publication, we identified 80, 87 and 90 essential genes for RKO, NCIH1299 and DLD1 cell lines, respectively (Supplementary Table 2). The union of the three consists 108 essential genes. If a gene is essential in more than one cell lines, we averaged the Z-scores for each sgRNA to increase signal-noise ratio.

### The ProTiler algorithm

ProTiler Takes three steps to identify CKHS regions: (1) weak signal removal and outlier adjustment; (2) Segmentation using TGUH method; (3) CKHS region calling from segments.

Approximately 1/3 of the sgRNAs with a PAM-appended target are inactive, corresponding to weak dropout effects^36, 44^. To remove weak dropout signals, each sgRNA data point is compared to its *k* neighbors to the left and *k* neighbors to the right. The data point is removed if the signal is weaker than 2/3 of left neighbors and 2/3 of right neighbors. We set *k*=5, corresponding to an average window span of ~30 AAs, the size of the smallest protein domain module. To adjust the outliers, we estimate the variation of noise for each protein, by applying Median Absolute Deviation (MAD) on the differences between consecutive sgRNA signal. For each data point *x*, we compare it to the median value of its neighbors within a sliding window of size 11. If *x* is larger than the median value by more than twice of MAD, *x* is marked to be an outlier and is adjusted to be median+2*MAD. The outliers below the median values are detected and adjusted in a similar way.

To segment the protein into regions corresponding to hyper- or hypo-sensitivity, ProTiler uses a Tail-Greedy Unbalanced Haar (TGUH) method, which decomposes noisy 1-D data and detects multiple change-points based on wavelet transformation^11^. Different from regular binary segmentation methods that adopt a ‘top-down’ strategy to search for segments, TGUH uses a ‘bottom-up’ strategy via a natural unary-binary tree, making it more accurate in recognizing small segments. In ProTiler, TGUH is implemented using R package library “breakfast”.

Suppose we have *n* segments, and the *i*th segment contains *k*_*i*_ data points (*i*=1, 2, …, n), we assigned a score, *s*_*i*_, to be the average of data points in the *i*th segment. The segments are sorted in ascending order of scores, such that *s*_1_ ≤ *s*_2_ ≤ ⋯ ≤ *s*_*n*_. Since a more negative value corresponds to a stronger dropout effect, the sorted list is in descending order of CRISPR knockout sensitivity. We iteratively assign the segments to the CKHS category. The pseudo-code of CKHS region calling is as follow.

~~~
BEGIN
assign the first segment to be CKHS
for *i* in 2 to *n*
    compute 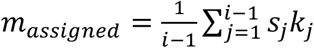
    compute 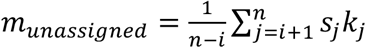
    if *s*_*i*_ < (*m*_*assigned*_ + *m*_*assigned*_)/2 assign the *i*th segment to be CKHS
    else
        break
merge adjacent CKHS segments in the protein into a single CKHS region
END
~~~

### Prediction of CKHS region

The protein features were extracted and encoded as follow:

- Domain annotation: Each AA is assigned 1 if it is within a Pfam domain, otherwise assigned 0.
- Conservation score: SIFT scores were computed for each amino acid, followed by a Gaussian kernel smoothing with bandwidth=10AA. The conservation scores were mean-centered for each protein.
- Secondary structure: The secondary structures were predicted using RaptorX and were mapped to each AA. We assigned a code of [1,0] for alpha helix, [0,1] for beta sheet, and [0,0] for unstructured.
- PTM: The annotations of phosphorylation, acetylation, methylation, and ubiquitination were mapped to the protein, followed by a Gaussian kernel smoothing with bandwidth=10AA.

A bagging SVM model was implemented for prediction^33^. The model contains 100 SVMs. In each iteration of bootstrapping, 5% of the AAs were randomly selected with replacement from the CKHS regions, and the same number of AAs were selected from the non-CKHS regions. The final prediction score is the average of the outputs from all SVMs. The SVMs were implemented using R-package library “e1071”.

To predict CKHS region in a proteome-wide scale, the bagging SVM model was trained using the CKHS regions identified in Munoz dataset, and was applied to all the CDS proteins. Predicted CKHS regions were further merged if their distance is less than 3 AAs. To reduce false positives, regions shorter than 10 AAs were discarded. In the analysis of Avana and GeCKO datasets, the sgRNA dropout effects were estimated using CERES^45^, and were averaged across the cell lines. 212 predefined core essential genes^34^ were used For Fig. 4h and Supplementary Fig. 5.

### Cell culture

The colon cancer cell line DLD-1 was obtained from ATCC (CCL-221). Cells were maintained in RPMI-1640 medium with 10% fetal bovine serum (FBS) and 1% Penicillin-Streptomycin. Medium was refreshed every 2-3 days. HEK293T cells were cultured with DMEM medium supplemented with 10% FBS and 1% Penicillin-Streptomycin.

### sgRNA design and nucleotide modification of the ectopic *SMARCB1*

The tiling sgRNA sequences of SMARCB1 were obtained from the previous publication^6^. The top three sgRNAs targeting SNF5_2 domain with most negative z-scores were chosen for further experiments. To ectopically expression of SMARCB1, the third bases of codons except the bases at PAM sites and methionine codon in the sgRNA targeting sequences were switched to other bases without coded amino acids changes. The tandem sequences coding Myc and 6xHis tags were appended at the 3’ terminus of modified *SMARCB1*. To generate truncated SMARCB1, the nucleotides coding the first 53 amino acids were removed and replaced by start codon on the basis of mutant full-length SMARCB1.

### Plasmid construction

Human mutant full-length and truncated *SMARCB1* were synthesized with modified nucleotides (Biomatik, USA). To construct overexpression plasmid, the mutant full-length and truncated fragments with Myc and His tags were respectively subcloned into pLVX-IRES-tdTomato vector (#631238, Clontech, USA) using restriction sites XbaI and BamHI. To knockout endogenous SMARCB1, sgRNA oligos were synthesized (VectorBuilder, USA) and cloned into lentiCRISPRv2 (#52961, Addgene) according to the protocol from Feng Zhang lab. The sgRNAs targeting *AAVS1* gene were used as the controls.

### Virus packaging and infection

HEK293T cells (4×10^6^) were seeded into 10 cm cell culture dishes one day before transfection in fresh medium. Before transfection, 4 µg target plasmid, 4 µg psPAX2 and 2 µg pMD2.G plasmids were added in 1 ml pre-warmed Opti-MEM medium (Gibco, # 31985062), and then mixed with 24 µl X-tremeGene HP DNA Transfection Reagent (Roche, #6366236001) at room temperature for 30 min. The mixture was dropwise added into each 10 cm dish containing HEK293T cells. Virus supernatant was collected 48 h after transfection, filtered through a 0.45 µm Acrodisc syringe filter, frozen in small volume and stored at −80°C until use. For infection, cells were seeded into 6-well plates with 5×10^5^ cells/well. After cells attached, lentivirus and 2 µl polybrene (Millipore, #TR-1003-G) were added with totally 2 ml medium in each well. Forty-eight hours after infection, cells were seeded into 10 cm dishes for Puromycin (2 µg/ml) selection. To determine multiplicity of infection (MOI), different volumes of lentivirus were used for infection. Cell survival rate was calculated after Puromycin selection.

### Knockout and ectopic expression of SMARCB1

To test the knockout effects of sgRNAs targeting SMARCB1, viral lentiCRISPRv2-AAVS1sg and lentiCRISPRv2-SMARCB1sg were used for infection, followed by 3-7 days of Puromycin selection. To ectopically express mutant full-length and truncated SMARCB1, DLD-1 cells were infected with lentivirus harboring pLVX-IRES-tdTomato, pLVX-IRES-tdTomato-mutFullSMARCB1 and pLVX-IRES-tdTomato-mutTruncSMARCB1, cultured for several days and then sorted by FACS, respectively. For knockout rescue experiment, the sorted cells were infected with viral lentiCRISPRv2-AAVS1sg and lentiCRISPRv2-SMARCB1sg for SMARCB1 knockout and subsequently selected by Puromycin.

### Western blot

The knockout and overexpression of SMARCB1 were verified by Western blot. Briefly, the cells sorted by FACS and one week after lentiCRISPRv2-SMARCB1sg transfection were collected for protein extraction. The cells with pLVX-IRES-tdTomato and lentiCRISPRv2-AAVS1sg were used as the controls. Proteins were separated by SDS-PAGE and transferred onto PVDF membranes by semi-dry transferring system (Bio-rad, USA). After blocking with 5% skimmed milk, membranes were incubated with primary antibodies of rabbit anti-SMARCB1 (#A301-087A-M, Bethyl), anti-His (#12698T, CST) and anti-Myc (2278T, CST) at the concentration of 1:2000 at 4°C overnight. After TBST washing, secondary anti-rabbit (#NA934, GE) and mouse (#NA931, GE) antibodies were used for incubation at 1:10000 for 1-2 h. The bands were visualized by ECL reagent (#NEL104001EA, PerkinElmer). β-actin was used as the loading control.

### Cell growth assay

After 3-7 days of Puromycin selection, the cells transfected with lentiCRISPRv2-SMARCB1sg were seeded into 96-well plates at the density of 1000 cells/well in 100 µl medium. Cell proliferation was determined by CellTiter-Glo assay (Promega, #G7572) daily according to the manufacturer’s instructions. Triple independent repeats were conducted.

## Data availability

The processed data are available in the Supplementary Tables of this article. The graphic views of CKHS profiles of 83 proteins are publicly available at https://figshare.com/s/27d52df30b2bcc7a038a.

## Software code availability

ProTiler was written in Python (version 2.7) and R package (version > 3.5.0), implemented as open source software downloadable from https://github.com/MDhewei/ProTiler-1.0.0.

## Acknowledgements

We thank Drs. Sharon Dent and Xiaodong Cheng for critical discussion, and Dr. Briana Dennehey for manuscript inspection. This work was supported by a CPRIT grant RR160097 (H.X.).

## Author contributions

W.H., L.Z., and H.X. conceptualize the study. W.H. developed the software. W.H., O.D.V., and J.D. performed computational analysis. L.Z., R.F., and E.B. performed biochemistry experiments. M.T.B, X.S., T.C., and B.B. helped data interpretation. H.X. supervised the project. All authors participate in writing the manuscript.

